# Mechanisms of spinophilin-dependent pancreas dysregulation underlying diabesity

**DOI:** 10.1101/2023.02.07.527495

**Authors:** Kaitlyn C. Stickel, Amber L. Mosley, Emma H. Doud, Teri L. Belecky-Adams, Anthony J. Baucum

## Abstract

**Objective:** Spinophilin is an F-actin binding and protein phosphatase 1 (PP1) targeting protein that acts as a scaffold of PP1 to its substrates. Spinophilin knockout (Spino^-/-^) mice have decreased fat mass, increased lean mass, and improved glucose tolerance, with no difference in feeding behaviors. While spinophilin is enriched in neurons, its roles in non-neuronal tissues, such as beta cells of the pancreatic islets, are unclear.

**Methods & Results:** We have corroborated and expanded upon previous studies to determine that Spino^-/-^ mice have decreased weight gain and improved glucose tolerance in two different models of obesity. Using proteomics and immunoblotting-based approaches we identified multiple putative spinophilin interacting proteins isolated from intact pancreas and observed increased interactions of spinophilin with exocrine, ribosomal, and cytoskeletal protein classes that mediate peptide hormone production, processing, and/or release in Lepr^db/db^ and/or high fat-fed (HFF) models of obesity. Moreover, loss of spinophilin specifically in pancreatic beta cells improved glucose tolerance without impacting body weight.

**Conclusion:** Our data further support a role for spinophilin in mediating pathophysiological changes in body weight and whole-body metabolism associated with obesity and provide the first evidence that spinophilin mediates obesity-dependent pancreatic dysfunction that leads to deficits in glucose homeostasis or diabesity.

## INTRODUCTION

Chronic obesity is associated with pathophysiological changes that predispose individuals to dysregulation of glucose uptake and pancreatic beta cell dysfunction that contribute to the development of Type 2 diabetes (T2D) (1). The presentation of diabetes due to obesity/overweight, termed ‘diabesity’, is a major public health concern (2, 3). Obesity can lead to peripheral insulin resistance that, if not compensated for by increased insulin secretion by the beta cells, will lead to dysregulated glucose uptake and hyperglycemia that defines T2D (4). To compensate for increases or decreases in peripheral (to the beta cell) insulin sensitivity, the beta cell can non-linearly decrease or increase insulin secretion in response to glucose (4). Moreover, high glucose stimulation or cellular depolarization with potassium chloride induces biphasic insulin secretion. In isolated islets, the initial phase lasts ∼10 minutes and the more prolonged phase ∼25 – 35 minutes (5). The first phase is thought to involve readily-releasable large dense core granules (LDCGs) that employ multiple membrane receptors, including G-protein coupled receptors (GPCRs), ion channels, and receptor tyrosine kinases along with downstream kinase activation and calcium-dependent insulin exocytosis to respond to glucose-stimulation (4). The second phase of insulin secretion is induced by movement of the reserve pool of LDCGs via cytoskeleton rearrangement (6). In addition to regulation of insulin by glucose-stimulated insulin secretion (GSIS), there is transcriptional, posttranscriptional, translational, and posttranslational regulation of insulin production and stability which can modulate the amount of insulin that is contained within LDCGs of the beta cells (7, 8, 9, 10, 11, 12). However, mechanisms directly linking obesity and diabetes together are poorly understood.

Spinophilin is a brain-enriched protein phosphatase 1 (PP1) targeting protein that is implicated in neuronal adaptations. Initial characterization of mice with global knockout (KO) of spinophilin (Spino^-/-^) found decreased body weight compared to wildtype (WT) mice (13). Moreover, more recent studies have found that Spino^-/-^ mice have improved glucose uptake and reduced weight gain, measures associated with improved insulin sensitivity and metabolic function (14, 15). While we and others have characterized the importance of spinophilin in synaptic signaling mechanisms in the brain (13, 16, 17, 18, 19, 20, 21, 22), its expression and role in non-neuronal tissues such as the pancreas is less clear.

Previous studies identified that 16–18-week-old chow-fed, male Spino^-/-^ mice had decreased fat mass, increased lean mass, and improved glucose tolerance (14). They proposed a beta cell signaling mechanism from *in vitro* studies via M3 muscarinic acetylcholine receptors (M3R) for spinophilin’s involvement in negatively regulating M3R signaling and first phase insulin release in MIN6 beta cells (14). Moreover, recent studies found significant differences in weight gain, glucose uptake, and insulin sensitivity only in male spinophilin knockout (KO) versus WT mice on an 8-week high fat diet, with no significant differences in the female population, and proposed a mechanism involving spinophilin signaling in adipose tissue (15, 23). However, pancreas-specific mechanisms by which loss of spinophilin improves metabolic parameters are unknown.

In this study, we found that loss of spinophilin attenuates weight gain in both male and female Lepr^db/db^ and high-fat diet fed (HFF) obese mice and improves glucose tolerance. Using unbiased proteomics approaches and targeted immunoblotting, we have found alterations in spinophilin interactions in the pancreas isolated from different obesity mouse models. Specifically, we identified overall increases in spinophilin protein interactions in the pancreas of HFF mice with proteins that are classically involved in signaling, protein translation, and cytoskeletal rearrangement in the beta cell, pathways that are all critical in insulin processing/release. Finally, we found that loss of spinophilin only in insulin producing beta cells improves glucose tolerance in a cohort of young, chow-fed, male mice. Overall, we found that while spinophilin decreases weight gain, it improves glucose tolerance via pancreatic beta cell-specific mechanisms, potentially via its interactions with multiple proteins that involved in hormone production, processing, and release. These data position spinophilin at multiple points within the pancreas and pancreatic beta cells to regulate diabesity.

## MATERIALS & METHODS

### Animals

See Supplemental Methods.

### Intraperitoneal Glucose Tolerance Test (IPGTT)

Fasting GTTs were performed at 6 and 10 weeks of age. Mice were fasted for 4 hours and an initial blood glucose reading was taken. Mice were injected intraperitoneally (I.P.) with glucose (2 g dextrose/ kg body weight) and blood glucose was measured 15-, 30-, 60-, 90-, and 120-minutes post-injection using blood glucose test strips & monitor (Alpha-Trak2, Zoetis, Inc, Parsippany, NJ).

### Intraperitoneal Insulin Tolerance Test (IPITT)

ITTs were performed at 8 weeks of age by fasting the mice short-term (4 hours). An initial blood glucose reading was taken prior to injection with 1 U/kg body weight of insulin (Humulin U-100, Eli Lilly and Co, Indianapolis, IN, catalog #4273850). An intraperitoneal (I.P.) injection of insulin was given to the mice after the fasting period, and blood glucose was monitored at 15-, 30-, 60-, 90-, and 120-minutes post-injection using blood glucose test strips & monitor (Alpha-Trak2).

### Immunoprecipitations, Immunoblotting, and Proteomics Studies

See supplemental methods for methodology and references

### Statistics

Analysis of curves was assessed by performing t-tests, one-way ANOVAs, two-way ANOVAs, three-way ANOVAs and appropriate post-hoc tests. A Grubbs’s test was performed to identify outliers in data. Statistical significance was set at p<0.05. Specific statistical tests and subsequent post-hoc tests with results are fully listed in **Table S1**. All analyses were performed in Prism (GraphPad). For all studies, a single animal is the experimental unit.

## RESULTS

### Loss of spinophilin attenuates weight gain in Lepr^db/db^ obese mice and mice on a HFF diet

Previous studies established that loss of spinophilin resulted in reduced weight gain in HFF & chow-fed, male spinophilin KO mice compared to WT mice (14, 15) while HFF spinophilin KO females had no difference in body weight. Male and female Spino^-/-^/Lepr^db/db^ and Spino^-/-^/Lepr^+/+^ mice gained significantly less weight than their corresponding Spino^+/+^ littermates (**Figures 1 A-B**). Spino^-/-^ male and female HFF mice also gained significantly less weight than Spino^+/+^ HFF littermates of the corresponding sex (**Figures 1 C-D**). Therefore, loss of spinophilin in male and female mice significantly reduces weight gain in both lean mice and in multiple obesity models when measuring long-term weight changes.

**Figure 1.**
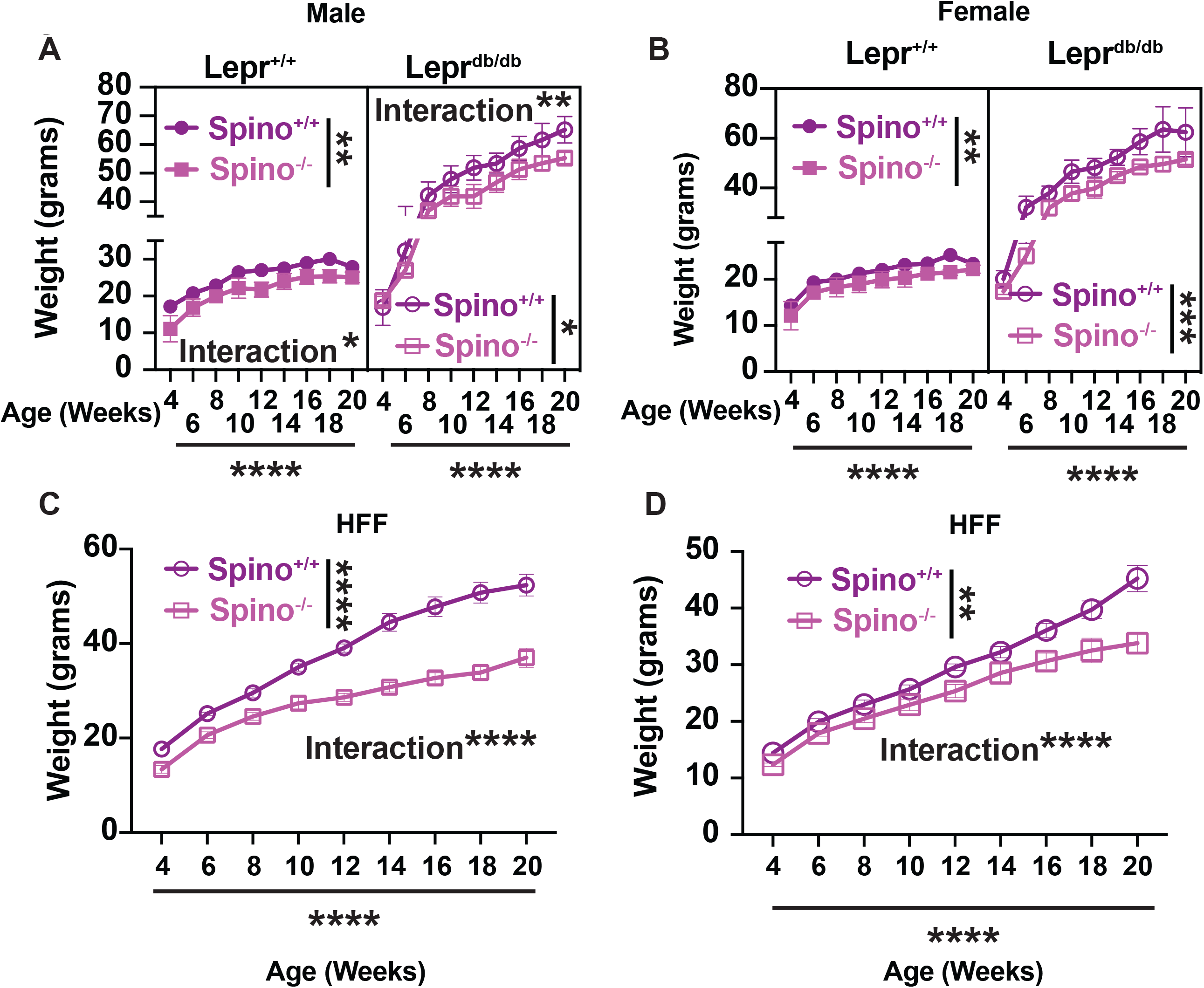
Loss of spinophilin significantly reduces weight gain in lean & two mouse models of obesity. **A, B**. Male (**A**) or female (**B**) Lepr^+/+^/Spino^+/+^, Lepr^db/db^/Spino^+/+^, Lepr^+/+^/Spino^-/-^, and Lepr^db/db^/Spino^-/-^ mice weights were taken bi-weekly from 4 to 20 weeks and plotted. A mixed effects 3-way ANOVA was performed initially followed by a two-way ANOVA to determine the effect of spinophilin genotype, age, and an interaction within the Lepr^+/+^ and Lepr^db/db^ genotypes individually. **C, D**. Male (C) and Female (D) Spino^+/+^/HFF and Spino^-/-^/HFF mice were weighed bi-weekly and plotted. For Two-Way ANOVAs, significant spinophilin genotype, time, and interaction (spinophilin genotype x time) are shown. N=4-12 for each age point. Data ± SEM are shown. *p<0.05, **p<0.01, ***p<0.001, ****p<0.001. All statistics are shown in Table S1.

### Loss of Spinophilin Improves GTT in Obese (HFF) Male & Female Mice

Previous studies concluded that loss of spinophilin improves GTT in male 16–18-week-old mice (14) and in 16-week-old male mice on high fat diet for 8 weeks, with no significant difference in HFF female mice (15). Our HFF male and female Spino^-/-^ mice were placed on a high fat diet starting at 4 weeks of age and GTTs were performed at 6 and 10 weeks of age. We found that loss of spinophilin in both male and female HFF mice had unique impacts on glucose tolerance at different ages. Specifically, male Spino^-/-^ HFF mice had no significant difference in glucose tolerance at 6 weeks of age (**Figures 2 A-B**) but weighed significantly less than WT mice (**Figure 2C**). At 10-weeks of age, male Spino^-/-^ mice had both significantly decreased GTT and significantly decreased body weights (**Figures 2 D-F**). At 6-weeks of age, female Spino^-/-^ mice had decreased GTT (**Figures 2 G-H**) and body weight (**Figure 2I**). At 10-weeks of age, there was a significant genotype effect on the GTT and a time x genotype interaction. However, there was only a trend for a decreased area under the curve for the GTT (**Figures 2 I-J**) and no significant difference in body weight between the two groups (**Figure 2 K**).

**Figure 2.**
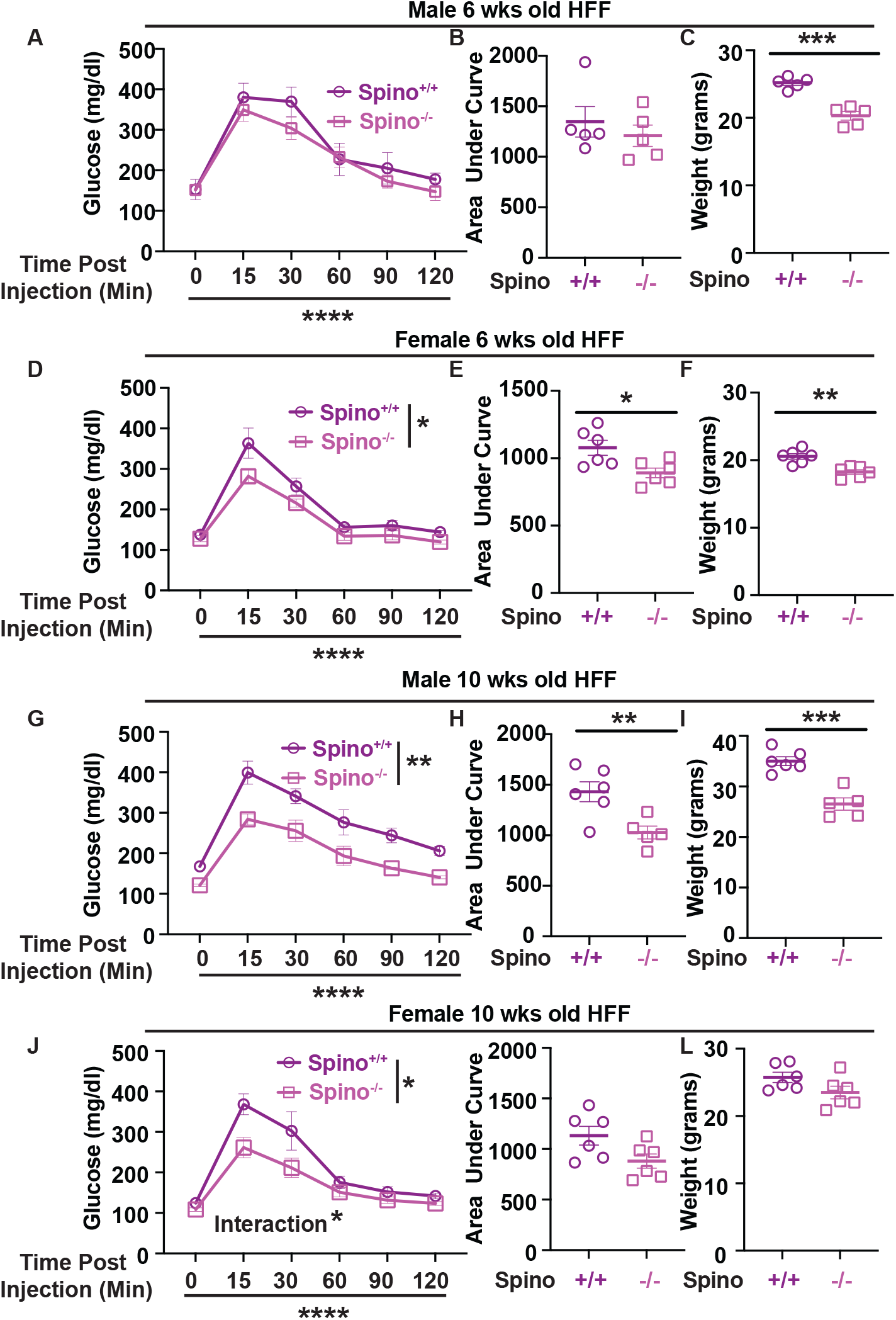
Spino^-/-^ mice have improved GTT in HFF male & female mice. **A**. Intraperitoneal glucose tolerance test (IPGTT) of HFF WT and spinophilin KO male mice at 6 weeks of age. **B**. Area under curve for the IPGTT from HFF WT and spinophilin KO male mice at 6 weeks of age. **C**. Weights from male HFF WT and spinophilin KO mice at 6 weeks of age. **D**. IPGTT of HFF WT and spinophilin KO female mice at 6 weeks of age. **E**. Area under curve for the IPGTT from HFF WT and spinophilin KO female mice at 6 weeks of age. **F**. Weights from female HFF WT and spinophilin KO mice at 6 weeks of age. **G**. IPGTT of HFF WT and spinophilin KO male mice at 10 weeks of age. **H**. Area under curve for the IPGTT from HFF WT and spinophilin KO male mice at 10 weeks of age. **I**. Weights from male HFF WT and spinophilin KO mice at 10 weeks of age. **J**. IPGTT of HFF WT and spinophilin KO female mice at 10 weeks of age. **K**. Area under curve for the IPGTT from HFF WT and spinophilin KO female mice at 10 weeks of age. **L**. Weights from female HFF WT and spinophilin KO mice at 10 weeks of age. Data are given with SEM. A 2-way repeated measures ANOVA (**A, D, G, J**) or unpaired t-tests (**B, C, E, F, H, I**) were performed. For Two-Way ANOVA, significant spinophilin genotype, time, and interaction (spinophilin genotype x time) are shown. N=5-6. *p<0.05, **p<0.01, ***p<0.001, ****p<0.001. All statistics are shown in Table S1.

### Obesity modulates spinophilin interactions in the pancreas

The weight gain and GTT data described above suggest loss of spinophilin may impact obesity and glucose tolerance independently. While previous studies using an immortalized mouse insulinoma cell line beta cell line (MIN-6) demonstrated a spinophilin-dependent regulation of M3 muscarinic receptor-dependent insulin-secretion (14), the role of spinophilin in vivo in the intact pancreas has not been probed. Using proteomics and immunoblotting-based approaches (**Figure 3A**), we identified multiple putative spinophilin interacting proteins from whole pancreas lysates isolated from WT and Lepr^db/db^ mice by immunoprecipitating for spinophilin and subjecting immunoprecipitates to in gel tryptic digestion followed by mass spectrometry (**Figure 3B-D; Tables 1, S2-S3**). We utilized STRING database (stringdb) (24) as part of the ELIXIR infrastructure to perform the Kyoto Encyclopedia of Genes and Genomes (KEGG) and Gene Ontology (GO) pathway analyses (**Tables S4-S7**) on all spinophilin interacting proteins identified in our proteomics experiment with a total spectral count of 4 or greater. We then clustered similar proteins together based on specific biological processes, cellular component/localization, and KEGG pathways (**Figure 3 B-D**). Spinophilin contains an F-actin binding domain and is important in cytoskeletal rearrangement in dendritic spines (16, 25). Here, we also identified that spinophilin interacts with different classes of myosins and actins in the pancreas that are important in cytoskeletal organization (26, 27, 28) (**Figure 3B**). Myosin-9 was identified to have the greatest difference between the number of spectral counts observed in the WT and Lepr^db/db^ mice. Interestingly, we observed multiple spinophilin interacting proteins involved in protein translation in the pancreas, including ribosomal proteins, heat shock proteins, and ER-chaperones that are upregulated in ER stress conditions such as BiP and protein disulfide isomerase (PDI) (**Figure 3C**). Moreover, we concluded spinophilin interacts with proteins classically identified in pancreatic secretion, insulin signaling, and protein digestion (**Figure 3D**).

**Table 1.** Full Pancreas Proteomics.

| **Table 1. Full**<br>**Pancreas**<br>**Proteomics** | **Goat Spinophilin Antibody** |  | **Rabbit Spinophilin Antibody** |  | **Total**<br>**Spectral**<br>**Counts** |
| --- | --- | --- | --- | --- | --- |
|  | **WT** | **Lepr<sup>db/db</sup>** | **WT** | **Lepr<sup>db/db</sup>** |  |
| Neurabin-2<br>(spinophilin) | **21** | **17** | **42** | **36** | **116** |
| Neurabin-1<br>(homolog-86%) | **1** | **0** | **10** | **0** | **11** |
| PP1α<br>catalytic<br>subunit | **0** | **1** | **3** | **4** | **8** |
| PP1γ<br>catalytic<br>subunit | **0** | **1** | **3** | **5** | **9** |
| BiP | **2** | **25** | **10** | **34** | **71** |
| Myosin-9 | **0** | **74** | **0** | **56** | **130** |

**Figure 3.**
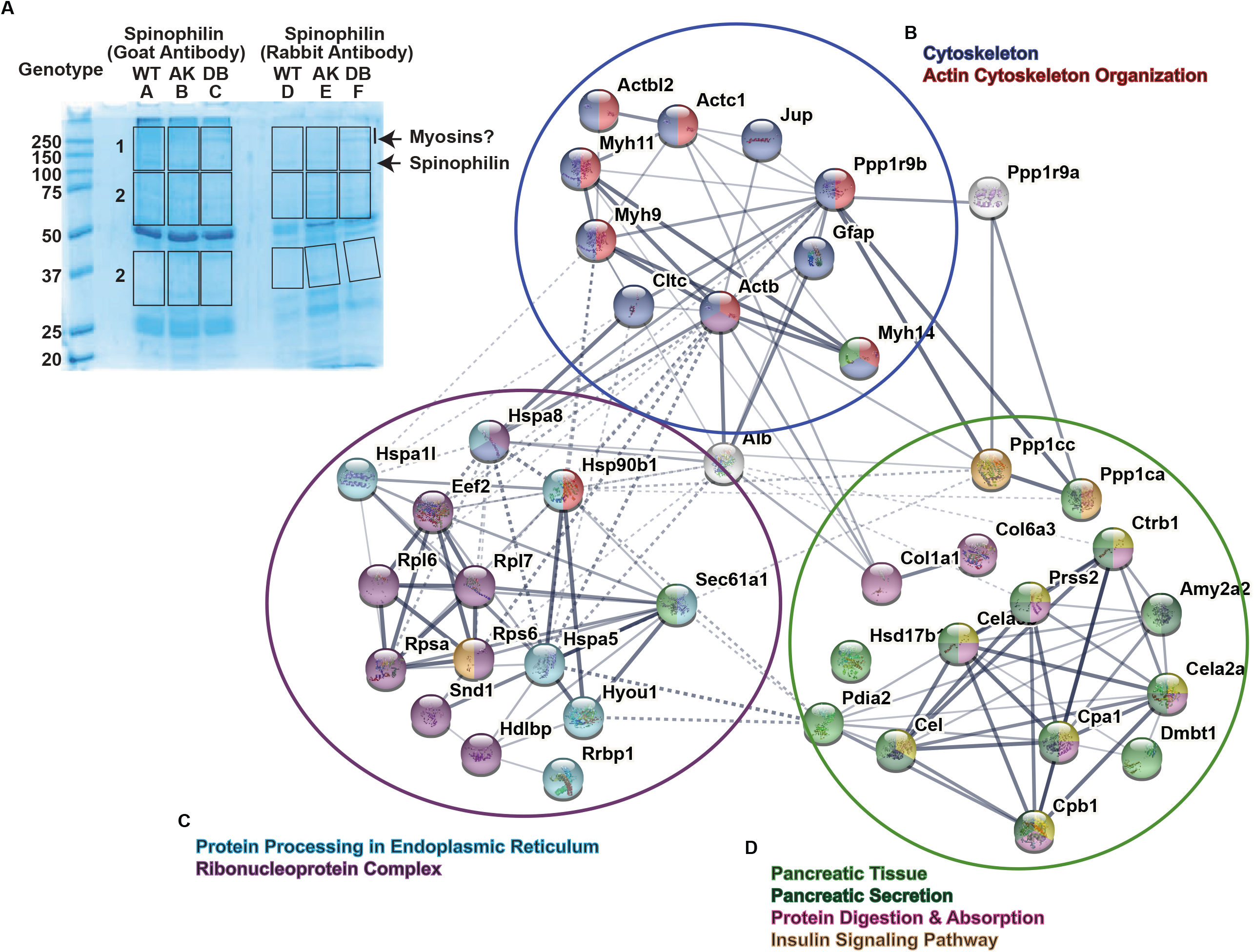
A. Coomassie Blue SDS-PAGE Gel Showing Spinophilin Immunoprecipitates. Spinophilin was immunoprecipitated from pancreas of a single 6-week-old WT and Lepr^db/db^ mice using two different spinophilin antibodies. Immunoprecipitates were separated by SDS-page, stained, and excised for MS-based analysis. 66 total interacting proteins were detected with 2 or more spectral counts (Supplemental Table 3) but for more targeted expression, we only considered proteins with spectral counts of 4 or above (Supplemental Table 2). **B-D**. Representation of KEGG pathway analysis from STRING input. Spinophilin interactors altered in obesity have been sorted into 3 clusters based on common biological processes. Interactors with a spectral count of 4 or more were included. **B**. Spinophilin interactors clustered into common cytoskeletal functions. **C**. Spinophilin interactors clustered into common biological functions involved in protein processing in the endoplasmic reticulum and ribonucleoprotein complexes. **D**. Spinophilin interactors clustered into common pancreatic functions, such as secretion, digestion, and insulin signaling. **Table 1**. Table representing top spinophilin interactors identified using MS-based analysis. This table shows spectral counts from goat and rabbit spinophilin antibodies individually, and total spectral counts represented in the last column.

### Spinophilin Interacting Proteins in different models of diabetes

We immunoprecipitated spinophilin from pancreatic lysates and immunoblotted for interacting proteins observed in our initial proteomics study, including neurabin, PP1α, and myosin-9 in different models of diabetes. It is important to note that many of the proteins were not detected in pancreatic lysate western blots, in part due to the low abundance of these proteins. However, spinophilin and associated proteins were readily detected when enriched by spinophilin immunoprecipitation. We observed a decreased PP1 and spinophilin association in the Type 2 diabetic mouse model (Lepr^db/db^), but not the Ins2^Akita^ model of Type 1 diabetes, compared to WT mice. We observed a significant decrease in the association of spinophilin with its homolog neurabin in Lepr^db/db^, but not Ins2^Akita^ mice, compared to WT mice. Moreover, we only observed a western blot band with spinophilin & myosin-9 co-immunoprecipitation in our Lepr^db/db^ mice, with no association in WT or Ins2^Akita^ pancreas (**Figure 4**). This further confirmed the obesity-dependent increase in myosin-9 spectral counts in spinophilin immunoprecipitates observed in our proteomics study (**Table 1**).

**Table 2.**
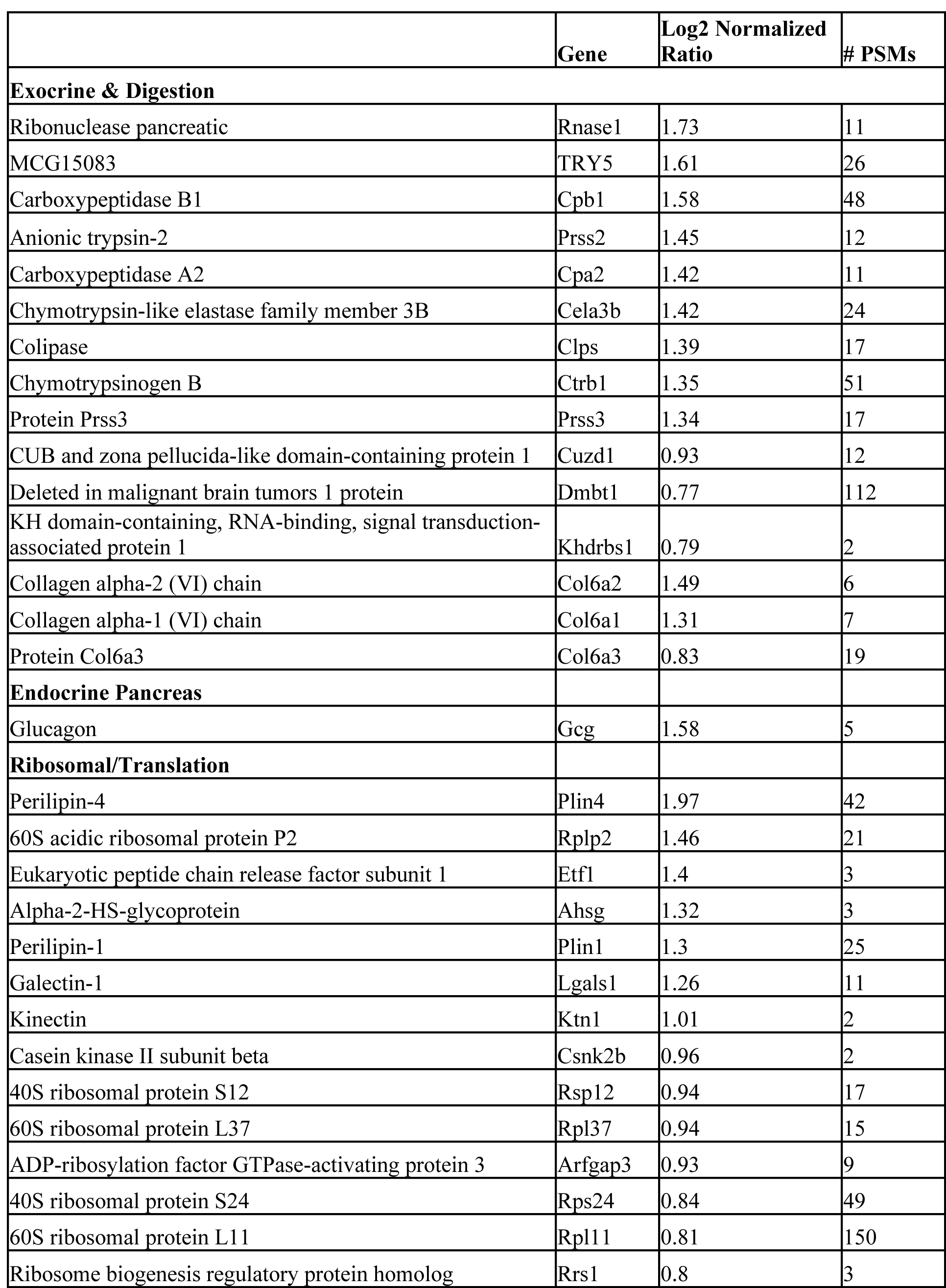
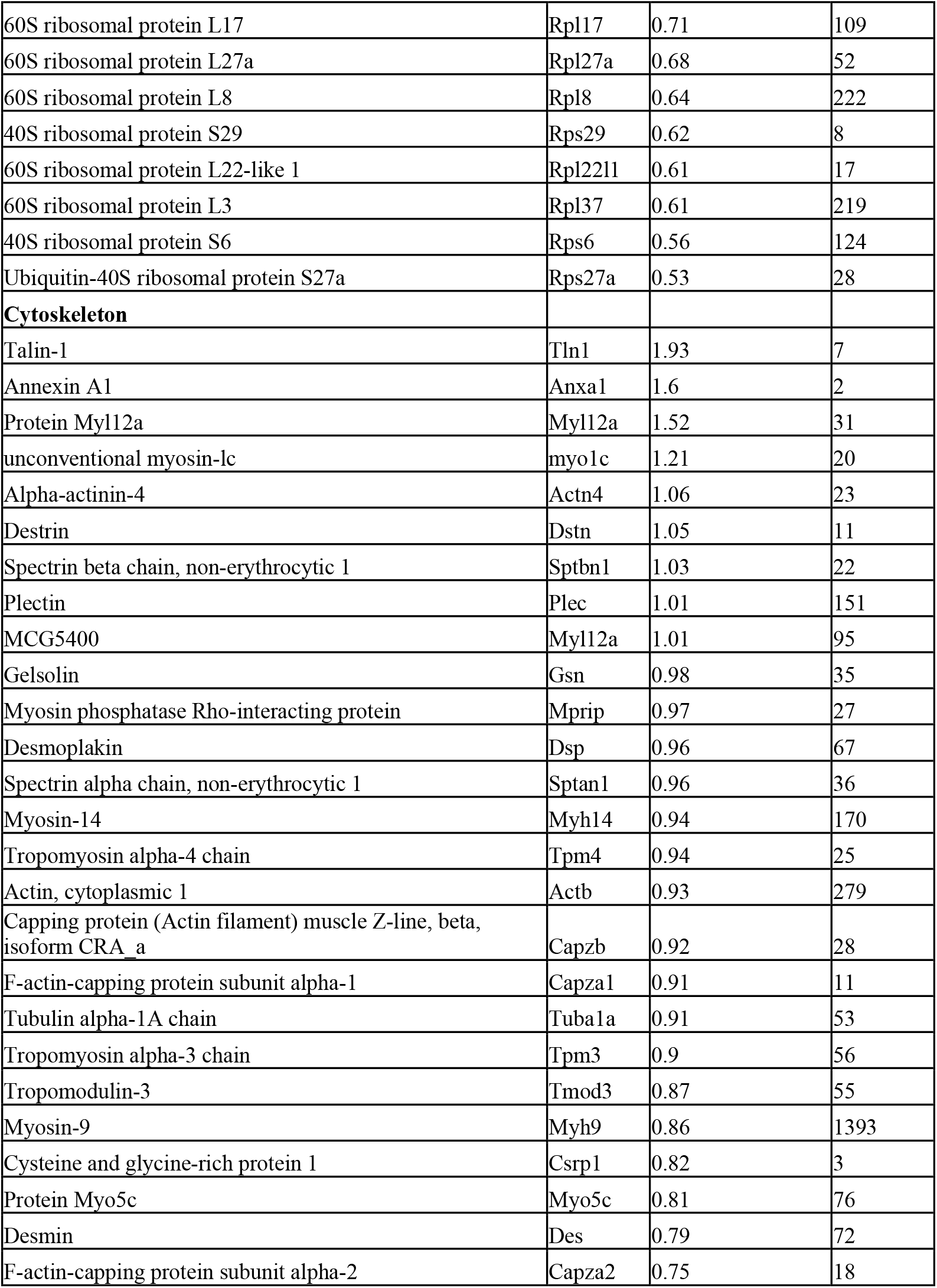
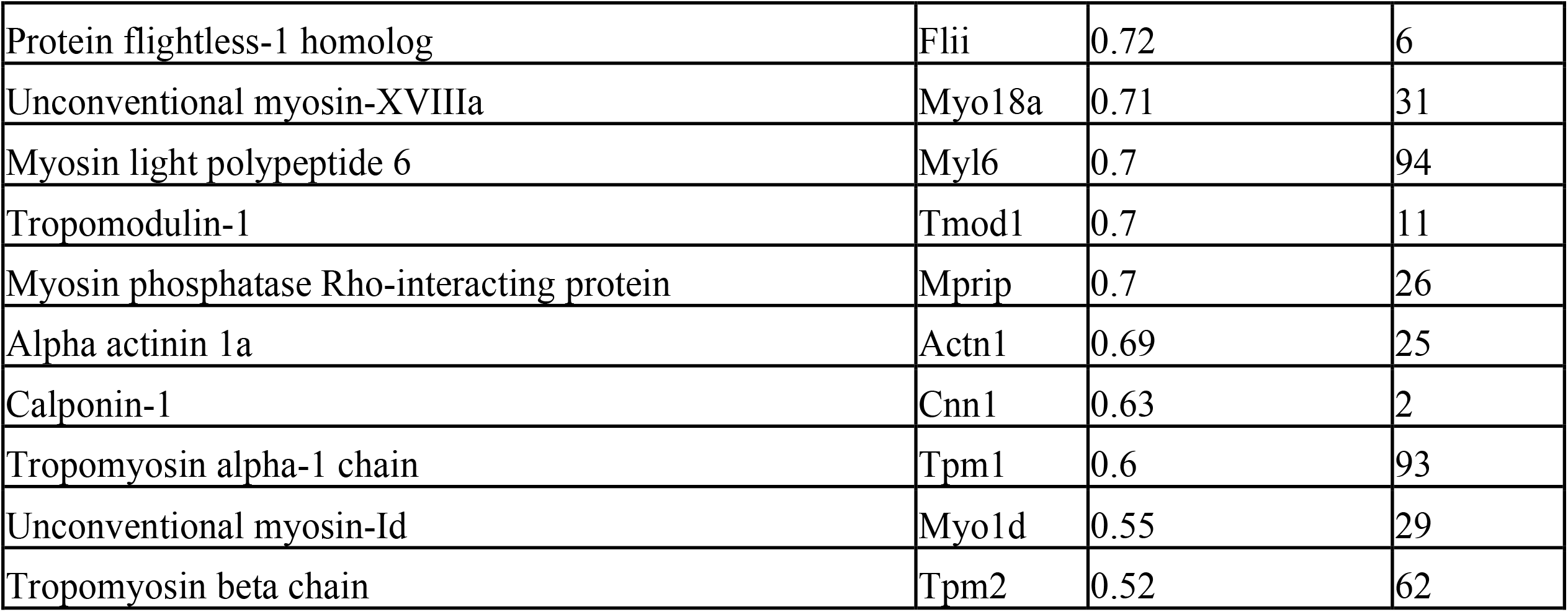
Spinophilin Interactors-Clusters.

**Figure 4.**
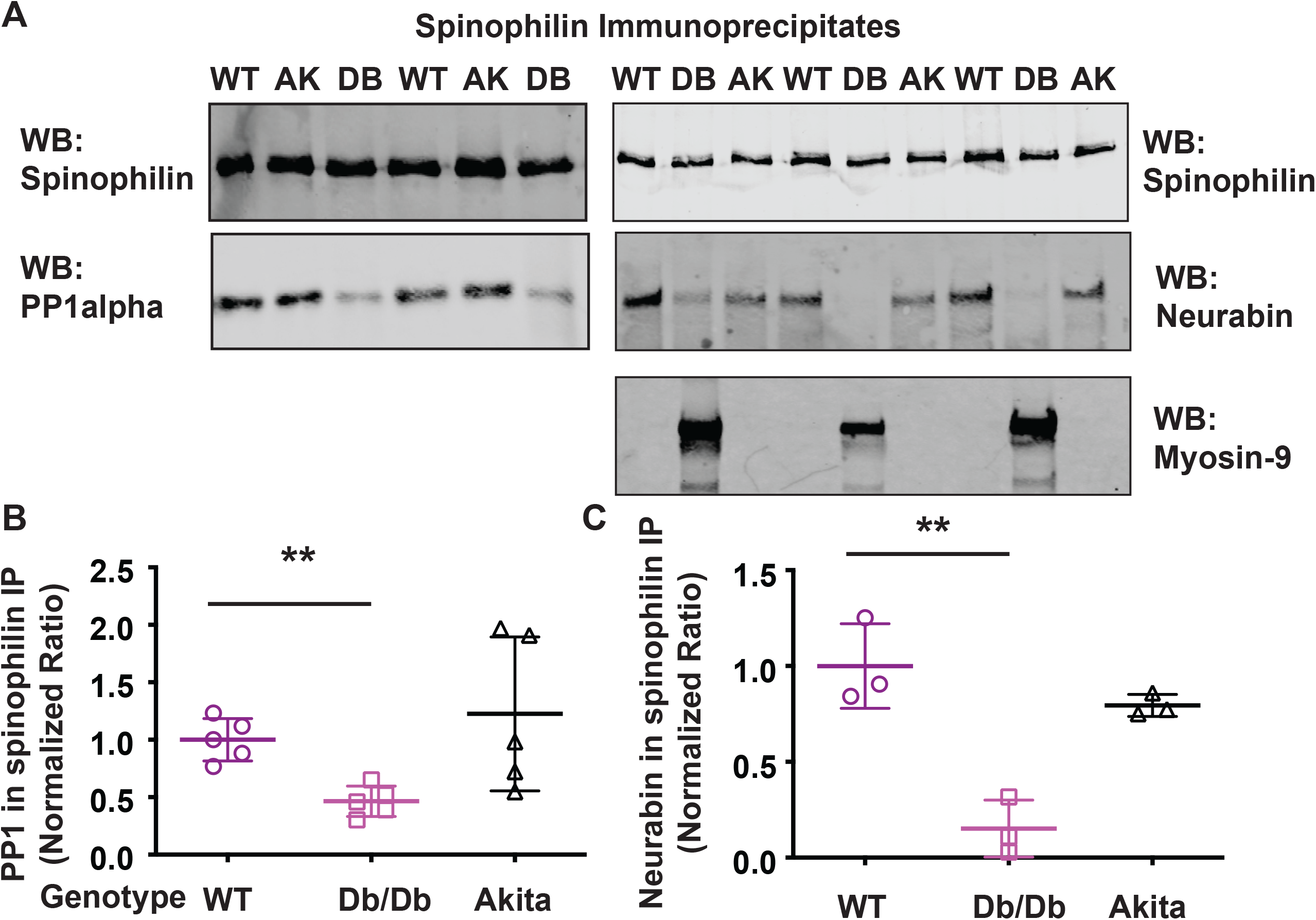
Spinophilin protein interactions with PP1, neurabin, and myosin-9 in wildtype, Ins2^Akita^, and Lepr^db/db^ mice. **A**. Spinophilin was immunoprecipitated from pancreas of adult WT, Ins2^Akita^ (AK), and Lepr^db/db^ (DB) mice. Immunoblotting for spinophilin, PP1α, myosin-9, and neurabin was performed. **B**. PP1α expression in spinophilin immunoprecipitates was normalized to spinophilin expression in the immunoprecipitate. A normalized ratio was plotted. N=3-5. **p<0.01. All statistics are shown in Table S1.

### TMT proteomics of HFF WT & Spinophilin KO mouse pancreas

To validate and quantify these interactions in another model of obesity and to determine specificity of the interactions with spinophilin, we immunoprecipitated spinophilin from the pancreas of WT lean male mice on standard chow and WT HFF obese male mice. Additionally, we utilized HFF spinophilin KO mice as a critical negative interaction control (18). Immunoprecipitates were analyzed using a ratiometrically quantitative tandem mass tag (TMT) proteomics experiment. To be considered for our specificity cutoff and to remove any contaminates or non-specific interactors, we filtered our samples with a WT/KO log2 fold-enrichment of 0.5 and then removed any protein with less than 2 unique peptides. We then normalized to the spinophilin abundance in the corresponding sample and listed interactors (**Table S8**). All proteins detected regardless of specificity are shown in **Table S9**.

Overall, we replicated specific increased interactions with myosin-9 (**Table 2**) and BiP but did not observe a quantitative change in PP1 interaction with spinophilin in this obesity model. However, BiP protein was detected equally in WT and spinophilin KO HFF mice, suggesting an obesity-induced increase in this protein, but a non-specific pulldown (**Tables S8, S9**). We performed STRING, KEGG, and GO pathway analyses of specific spinophilin interacting proteins that also had an increased interaction (log_2_-fold change of ≥ 0.5in HFF vs. lean) (**Tables S10-S13**). We observed multiple proteins important in cytoskeletal organization (**Figure 5A**), translation (**Figure 5B**), and pancreatic secretion (**Figure 5C**).

**Figure 5.**
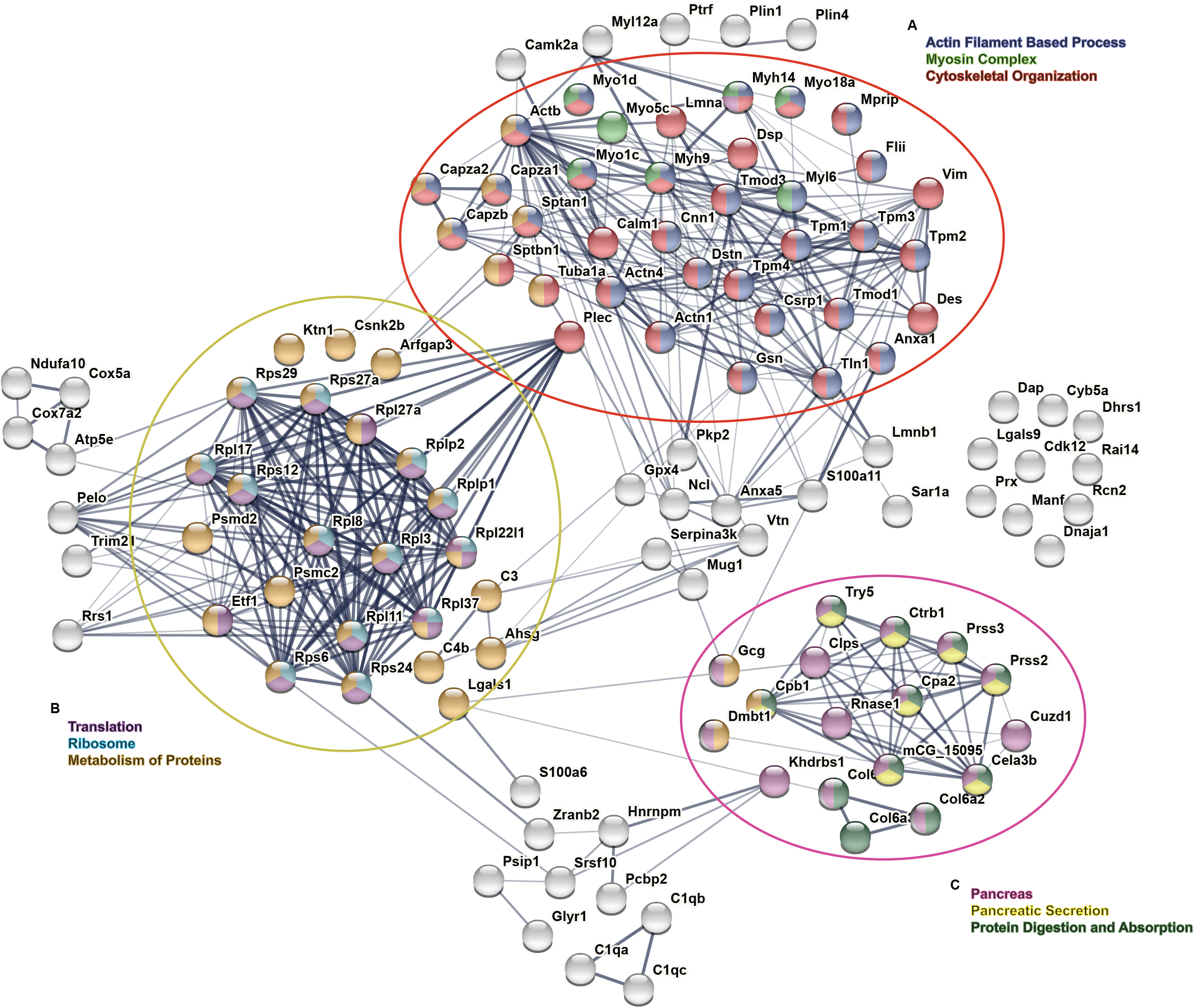
Visual representation of KEGG pathway analysis from STRING input. Spinophilin was immunoprecipitated from the pancreas of 3 WT lean male mice on standard chow, 3 WT HFF obese male mice, and 3 HFF spinophilin KO mice as a control. Immunoprecipitates were subjected to a tandem mass tag proteomics experiment. We then filtered our samples with a WT/KO log_2_fold-enrichment of ≥0.5, removed any protein with less than 2 unique peptides, and normalized to the spinophilin abundance in the corresponding sample. **A**. Top spinophilin interactors considered to have an obesity-dependent increase (log_2_-fold change) clustered together based on common functions in cytoskeletal organization. **B**. Top spinophilin interactors increased in obesity clustered into common biological functions, such as translation, metabolism of proteins, and ribosomes. **C**. Top spinophilin interactors increased in obesity clustered into common biological expression & function, including pancreas, pancreatic secretion, and digestion. A list of all spinophilin interactors in these clusters are provided in Table 2.

### Spinophilin in islets and pancreatic beta-cell specific spinophilin knockout

Overall, the proteomics data suggest that obesity modulates spinophilin protein-protein interactions within the pancreas, implicating novel roles for spinophilin in neurohormone production and release. To confirm a role for spinophilin in pancreatic beta cells *in vivo*, we isolated islets from mouse pancreas using a collagenase/protease solution and immunoprecipitated spinophilin. We were able to detect immunoprecipitated spinophilin in the isolated islets of mice using western blot (**Figure 6A**), indicating spinophilin is expressed in the highly secretory endocrine cells of the pancreas. Spinophilin is expressed in islets and loss of spinophilin globally improves pancreatic metabolic parameters such as glucose tolerance. To detail if loss of spinophilin specifically in pancreatic beta cells impacts GTT, we utilized our recently described conditional spinophilin knockout mice (Spino^fl/fl^) (29) and crossed them with Ins1Cre mice to knockout spinophilin specifically in pancreatic beta cells (Spino^ΔIns^). Spino^ΔIns^ mice, compared to control (Spino^fl/fl^ and Ins1Cre), had improved glucose tolerance at 6 and 10 weeks (**Figure 6 B, C, H, I**) with no change in body weight (**Figure 6 D, G, J**). There were no differences detected in the AUC between the two control groups (**Figure S1**), so we pooled the control area under the curve and weight data. When evaluating insulin tolerance, Spino^ΔIns^ mice had no deficits in insulin tolerance tests when data were not normalized to baseline (**Figure S2**). However, when normalized to baseline, Spino^ΔIns^ mice had significantly less glucose uptake upon insulin injection (**Figure 6 E-F**).

**Figure 6.**
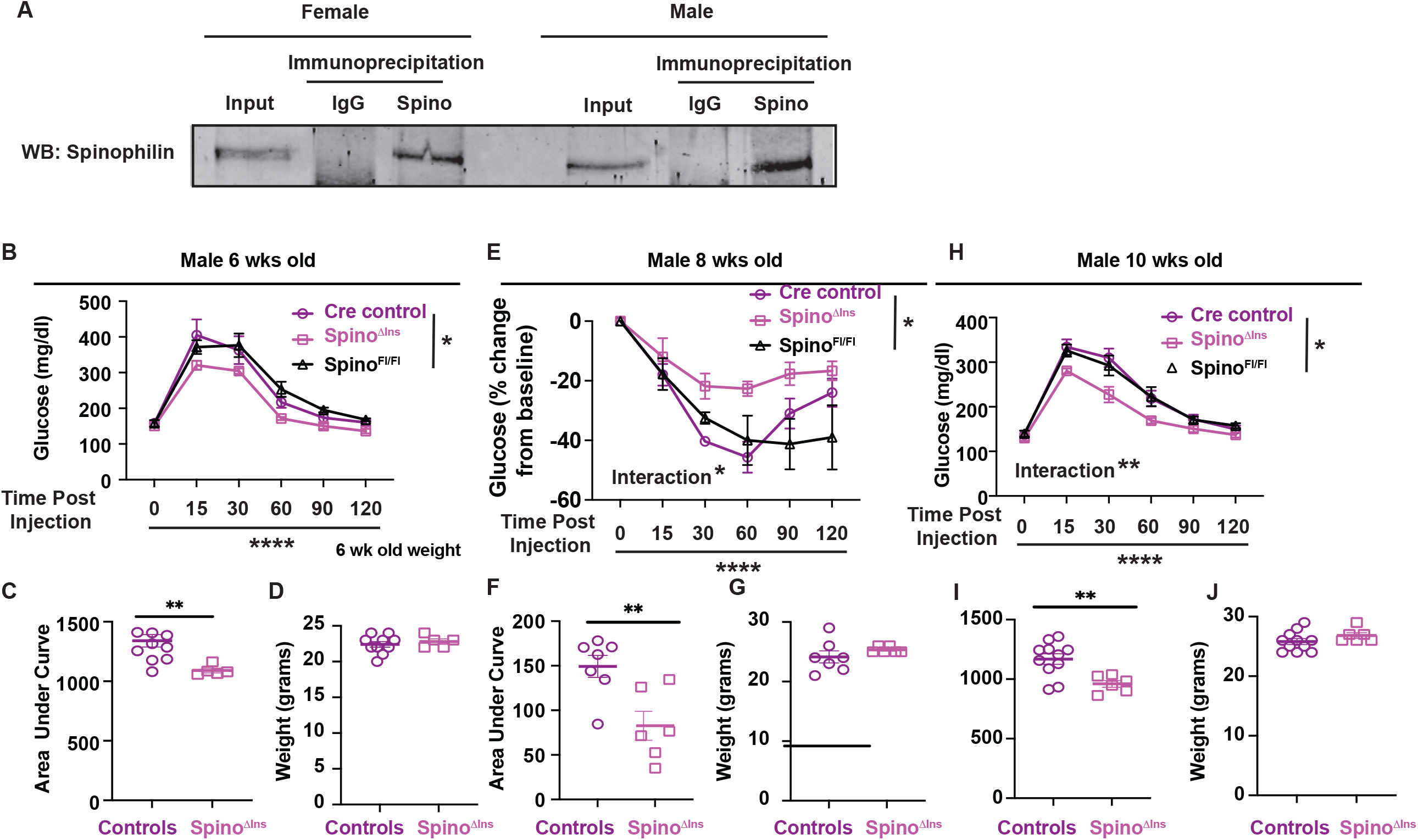
Loss of spinophilin, specifically in beta cells, improves glucose tolerance and reduces insulin tolerance. **A**. Spinophilin western blot of isolated islets from lysates or control IgG or spinophilin immunoprecipitates from a single male or female WT mouse. **B**. GTT of 6-week-old beta cell-specific spinophilin KO mice (Spino^ΔIns^) with Cre and flox controls. **C**. Area under curve for GTT of 6-week-old Spino^ΔIns^. **D**. Weights for 6-week-old Spino^ΔIns^ & controls. **E**. ITT (normalized to baseline) of 8-week-old Spino^ΔIns^ & controls. **F**. Area under curve for ITT (normalized to baseline) of 8-week-old Spino^ΔIns^ & controls. **G**. Weights for 8-week-old Spino^ΔIns^ & controls. **H**. GTT of 10-week-old Spino^ΔIns^ & controls. **I**. Area under curve for GTT of 10-week-old Spino^ΔIns^. **J**. Weights for 10-week-old Spino^ΔIns^ & controls. T-tests or Two Way ANOVAs (comparing genotype, time, or a genotype x time interaction) were performed. A Grubbs’s test was performed and identified two outliers, one 10 week old cre control and one 6 week old Spino^ΔIns^ mouse, which are not included in this data. N=3-7. *p<0.05, **p<0.01, ****p<0.001. All statistics are shown in Table S1.

## DISCUSSION

Two previous studies determined that Spino^-/-^ mice had lower body weight and improved glucose tolerance, but these changes were only tested or observed in male mice (13, 14, 15). Herein, we found that global loss of spinophilin in two mouse models of obesity, Lepr^db/db^ and HFF-induced obesity, decreased weight gain and improved glucose tolerance in both sexes. A previous study showed no difference in weights of HFF female mice; however, they started mice on diet at an older age (8 weeks vs. 4 weeks) and measured weights for 8 weeks compared to our 16 week evaluation (15). Therefore, earlier exposure or longer diet may impact female mice in addition to male mice. This is particularly true of the later ages as the difference between weights is greater at the later time points in the female mice.

Recent studies suggested changes in adipose tissue physiology may underlie improvements in metabolic parameters (15, 23). Interestingly, our observations that loss of spinophilin in Lepr^db/db^ mice decreased body weights in male and female mice to a similar extent suggests that additional mechanisms not associated with adipose tissue dysregulation of leptin hormone, at least, may be playing a role. Moreover, our data suggest that there may be a disconnect between weight gain and improved glucose tolerance. Therefore, we evaluated the impact of a high-fat diet on spinophilin in the pancreas. Using control and Lepr^db/db^ mice, we immunoprecipitated spinophilin from whole pancreas and performed a “GelC-MS” proteomics approach where we excised spinophilin immunoprecipitates from a Coomassie-stained gel. We detected multiple known interactors that we have previously observed in brain tissue (PP1, neurabin, and myosin protein) and putative novel (BiP and PDI) proteins. Overall, we identified proteins associated with actin cytoskeleton organization, ER protein processing, and pancreatic secretion. Using spectral counting, we found that HFF increased the interaction of multiple proteins with spinophilin, including myosin-9 and BiP, and decreased neurabin spectral counts in the Lepr^db/db^ mice compared to control mice. We validated the increased interaction with myosin-9 and the decreased interaction with neurabin. These data suggest that obesity impacts spinophilin interactions in the pancreas.

To follow-up these studies and further validate these changes, we compared spinophilin interactions from chow and HFF WT mice using a more advanced and ratiometrically quantitative TMT proteomics approach of the spinophilin immunoprecipitates. This approach enhances quantitation and increases the number of proteins detected as it does not require excision from a gel. In addition, these studies were performed on a more advanced, Orbitrap Eclipse Tribrid mass spectrometer. To further probe obesity effects on spinophilin interactions, we utilized a global, HFF spinophilin KO mouse to limit non-specific interactions. While we observed a greater total number of interacting proteins using this approach, there was overlap in the pathways that we detected when compared with the original Gel-C MS approach, including cytoskeletal proteins, translation proteins, and pancreatic secretion. Some proteins, such as BiP were not quantitatively higher in the HFF WT compared to spinophilin KO IP, suggesting this protein is a non-specific interactor with the spinophilin antibody or beads, but it may be generally upregulated under obese conditions. This experiment recapitulated the increased association of spinophilin with myosin-9 in obese mice. Myosins tend to be “sticky” when it comes to (immunoprecipitation and) mass spectrometry (30); however, it met our specificity and fold-change requirement cut-off, suggesting that this is a specific spinophilin interactor.

In addition to myosin-9, we observed increased spinophilin protein interactions in obesity with additional proteins involved in regulating actin dynamics, including: other myosin and myosin associated proteins, F-actin capping proteins, and actin proteins. Cytoskeletal proteins, such as class II & V myosins (28), have important roles in cytoskeletal rearrangement (27) and dense core vesicle transportation (31) that alter second phase GSIS in impaired beta cells, a long-term insulin secretion that requires movement of insulin granules to the membrane for release. Cytoskeletal rearrangement is known to contribute to the second phase of insulin secretion (27) and spinophilin is known to promote F-actin bundling via its F-actin binding domain (32). Therefore, it is possible that spinophilin may have specific impacts on the second phase of insulin secretion by enhancing the movement of the reserve pool of insulin-containing dense core vesicles and future studies will be required to detail if beta cell specific loss of spinophilin can impact phasic GSIS release either basally or following HFF.

We found an obesity induced increase in spinophilin interactions with proteins involved in protein digestion and absorption in the pancreas, such as trypsin, chymotrypsin, and collagenase. These proteins are classically associated with exocrine pancreas function; however, they also play a role in insulin processing (33) and future studies need to determine how spinophilin may modulate their specific functions.

We also observed several proteins involved in translation and protein metabolism, including ribosomal subunits and complement proteins. These proteins may be important in insulin processing and GPCR signaling (34). Spinophilin and GPCR signaling has been heavily investigated in the brain, but outside of the M3 muscarinic receptor (14), how spinophilin modulates GPCRs in the beta cells is unclear. We also identified multiple 40S and 60S ribosomal subunits that had an obesity-dependent increased interaction with spinophilin. Both subunits must be present for functional translation (35). However, ribosomal protein-deficient cells have impaired insulin signaling (36). An additional class of proteins that had an increased interaction with spinophilin was the perilipins, which protect the beta cell from another form of stress in T2D known as lipotoxicity (37). Perlipin-2 has previously been shown to regulate insulin secretion (38); however, the function of perilipin-1 and 4 detected here are less well-known. Future perifusion studies of islets from spinophilin knock out mice (global or cell type-specific) will be required to completely delineate spinophilin’s potential role in regulating phasic insulin secretion.

We observed multiple obesity-dependent changes in spinophilin interactions in the pancreas; however, whether beta cell specific loss of spinophilin function improves metabolic profiles was unclear. To address this, we generated and validated a Spino^ΔIns^ mouse. Interestingly, loss of spinophilin specifically in beta cells had no effect on weight gain but did improve glucose tolerance, at least in male mice, at 2 different ages. Future studies will need to detail if this effect is also present in female mice and if loss of spinophilin specifically in beta cells can rescue impairments in HFF-induced glucose intolerance. We observed less glucose uptake in response to insulin in Spino^ΔIns^ mice, contrasting previous studies that observed improved insulin-dependent glucose uptake (14, 15, 23). This difference may be due to different roles of spinophilin in islets vs peripheral tissues that take up glucose. For instance, given that beta cell type-specific loss improves glucose tolerance, basally, Spino^ΔIns^ mice may have greater circulating insulin which could occlude an effect of exogenous insulin. This would also contrasts with whole-body spinophilin KO mice which have less circulating insulin (14, 15) and therefore is an exciting avenue for future study. Our data are the first to measure metabolic parameters in our previously validated conditional spinophilin KO mice (29) and our current study demonstrates the need for cell type-specific approaches to fully understand spinophilin-dependent regulation of metabolism and cellular function.

Although we find here that levels of spinophilin expression in peripheral tissues impacts metabolic parameters, further work is needed to understand the signaling mechanisms by which spinophilin improves glucose tolerance and may regulate insulin secretion. As PP1 is highly promiscuous, modulating PP1 interacting proteins, such as spinophilin, offers a potentially more targeted approach to modulate this pathway. For instance, the FDA approved drug, Guanabenz, can act to modulate eukaryotic translation initiation factor 2 alpha signaling by modulating the GADD34-PP1 complex, demonstrating potential for altering PP1 activity by targeting regulatory proteins of the phosphatase (39). Therefore, future studies modulating inhibition of spinophilin-PP1 complex, or other spinophilin protein interactions could be a novel therapeutic approach for metabolic disorders related to obesity.

## Data availability

All TMT mass spectrometry data has been reposited to MassIVE ID: MSV000091159.

## ACKNOWLEDGMENTS

We thank Dr. Lisa Jones for use of the Q-exactive mass spectrometer and help with data analysis of the GelC-MS data. The TMT-mass spectrometry work was done by the Indiana University School of Medicine Center for Proteome Analysis. Acquisition of the IUSM Center for Proteome Analysis instrumentation used for this project was provided by the Indiana University Precision Health Initiative. We acknowledge and thank Wanda Filipiak & Galina Gavrilina for embryo injections for the initial generation of Spino^Fl/Fl^ mice as well as the entire excellent Transgenic Animal Model Core (in particular, Anna LaForest, Elizabeth Hughes, Corey Ziebell, and Dr. Thomas Saunders) and the University of Michigan’s Biomedical Research Core Facilities for their generation of these mice. We acknowledge Emily Claeboe for lab support and appreciate the feedback from all members of the Baucum laboratory on this project.

## References

1. Leitner DR, Fruhbeck G, Yumuk V, Schindler K, Micic D, Woodward E, et al. Obesity and Type 2 Diabetes: Two Diseases with a Need for Combined Treatment Strategies - EASO Can Lead the Way. Obes Facts 2017;10: 483–492.

2. Shafrir E, Ziv E, Kalman R. Nutritionally induced diabetes in desert rodents as models of type 2 diabetes: Acomys cahirinus (spiny mice) and Psammomys obesus (desert gerbil). ILAR J 2006;47: 212–224.

3. Farag YM, Gaballa MR. Diabesity: an overview of a rising epidemic. Nephrol Dial Transplant 2011;26: 28–35.

4. Kahn SE, Hull RL, Utzschneider KM. Mechanisms linking obesity to insulin resistance and type 2 diabetes. Nature 2006;444: 840–846.

5. Alcazar O, Buchwald P. Concentration-Dependency and Time Profile of Insulin Secretion: Dynamic Perifusion Studies With Human and Murine Islets. Front Endocrinol (Lausanne) 2019;10: 680.

6. Wang Z, Thurmond DC. Mechanisms of biphasic insulin-granule exocytosis - roles of the cytoskeleton, small GTPases and SNARE proteins. J Cell Sci 2009;122: 893–903.

7. Fu Z, Gilbert ER, Liu D. Regulation of insulin synthesis and secretion and pancreatic Beta-cell dysfunction in diabetes. Curr Diabetes Rev 2013;9: 25–53.

8. Lee EK, Gorospe M. Minireview: posttranscriptional regulation of the insulin and insulin-like growth factor systems. Endocrinology 2010;151: 1403–1408.

9. Kulkarni SD, Muralidharan B, Panda AC, Bakthavachalu B, Vindu A, Seshadri V. Glucose-stimulated translation regulation of insulin by the 5’ UTR-binding proteins. J Biol Chem 2011;286: 14146–14156.

10. Welsh M, Scherberg N, Gilmore R, Steiner DF. Translational control of insulin biosynthesis. Evidence for regulation of elongation, initiation and signal-recognition-particle-mediated translational arrest by glucose. Biochem J 1986;235: 459–467.

11. Wicksteed B, Alarcon C, Briaud I, Lingohr MK, Rhodes CJ. Glucose-induced translational control of proinsulin biosynthesis is proportional to preproinsulin mRNA levels in islet beta-cells but not regulated via a positive feedback of secreted insulin. J Biol Chem 2003;278: 42080–42090.

12. Sacco F, Humphrey SJ, Cox J, Mischnik M, Schulte A, Klabunde T, et al. Glucose-regulated and drug-perturbed phosphoproteome reveals molecular mechanisms controlling insulin secretion. Nat Commun 2016;7: 13250.

13. Feng J, Yan Z, Ferreira A, Tomizawa K, Liauw JA, Zhuo M, et al. Spinophilin regulates the formation and function of dendritic spines. Proc Natl Acad Sci U S A 2000;97: 9287–9292.

14. Ruiz de Azua I, Nakajima K, Rossi M, Cui Y, Jou W, Gavrilova O, et al. Spinophilin as a novel regulator of M3 muscarinic receptor-mediated insulin release in vitro and in vivo. FASEB J 2012;26: 4275–4286.

15. Zhang Y, Song L, Dong H, Kim DS, Sun Z, Boger H, et al. Spinophilin-deficient mice are protected from diet-induced obesity and insulin resistance. Am J Physiol Endocrinol Metab 2020;319: E354–E362.

16. Allen PB, Ouimet CC, Greengard P. Spinophilin, a novel protein phosphatase 1 binding protein localized to dendritic spines. Proc Natl Acad Sci U S A 1997;94: 9956–9961.

17. Allen PB, Zachariou V, Svenningsson P, Lepore AC, Centonze D, Costa C, et al. Distinct roles for spinophilin and neurabin in dopamine-mediated plasticity. Neuroscience 2006;140: 897–911.

18. Baucum AJ, 2nd, Jalan-Sakrikar N, Jiao Y, Gustin RM, Carmody LC, Tabb DL, et al. Identification and validation of novel spinophilin-associated proteins in rodent striatum using an enhanced ex vivo shotgun proteomics approach. Mol Cell Proteomics 2010;9: 1243–1259.

19. Edler MC, Salek AB, Watkins DS, Kaur H, Morris CW, Yamamoto BK, et al. Mechanisms Regulating the Association of Protein Phosphatase 1 with Spinophilin and Neurabin. ACS Chem Neurosci 2018;9: 2701–2712.

20. Morris CW, Watkins DS, Salek AB, Edler MC, Baucum AJ, 2nd. The association of spinophilin with disks large-associated protein 3 (SAPAP3) is regulated by metabotropic glutamate receptor (mGluR) 5. Mol Cell Neurosci 2018;90: 60–69.

21. Salek AB, Edler MC, McBride JP, Baucum AJ, 2nd. Spinophilin regulates phosphorylation and interactions of the GluN2B subunit of the N-methyl-d-aspartate receptor. J Neurochem 2019;151: 185–203.

22. Watkins DS, True JD, Mosley AL, Baucum AJ, 2nd. Proteomic Analysis of the Spinophilin Interactome in Rodent Striatum Following Psychostimulant Sensitization. Proteomes 2018;6.

23. Gou W, Wei H, Swaby L, Green E, Wang H. Deletion of Spinophilin Promotes White Adipocyte Browning. Pharmaceuticals 2023;16: 91.

24. Szklarczyk D, Gable AL, Nastou KC, Lyon D, Kirsch R, Pyysalo S, et al. The STRING database in 2021: customizable protein-protein networks, and functional characterization of user-uploaded gene/measurement sets. Nucleic Acids Res 2021;49: D605–D612.

25. Grossman SD, Hsieh-Wilson LC, Allen PB, Nairn AC, Greengard P. The actin-binding domain of spinophilin is necessary and sufficient for targeting to dendritic spines. Neuromolecular Med 2002;2: 61–69.

26. Kalwat MA, Thurmond DC. Signaling mechanisms of glucose-induced F-actin remodeling in pancreatic islet beta cells. Exp Mol Med 2013;45: e37.

27. Arous C, Rondas D, Halban PA. Non-muscle myosin IIA is involved in focal adhesion and actin remodelling controlling glucose-stimulated insulin secretion. Diabetologia 2013;56: 792–802.

28. Brito C, Sousa S. Non-Muscle Myosin 2A (NM2A): Structure, Regulation and Function. Cells 2020;9.

29. Morris CW, Watkins DS, Shah NR, Pennington T, Hens B, Qi G, et al. Spinophilin limits metabotropic glutamate receptor 5 scaffolding to the postsynaptic density and cell type-specifically mediates excessive grooming. Biological psychiatry.

30. Mellacheruvu D, Wright Z, Couzens AL, Lambert JP, St-Denis NA, Li T, et al. The CRAPome: a contaminant repository for affinity purification-mass spectrometry data. Nat Methods 2013;10: 730–736.

31. Varadi A, Tsuboi T, Rutter GA. Myosin Va transports dense core secretory vesicles in pancreatic MIN6 beta-cells. Mol Biol Cell 2005;16: 2670–2680.

32. Satoh A, Nakanishi H, Obaishi H, Wada M, Takahashi K, Satoh K, et al. Neurabin-II/spinophilin. An actin filament-binding protein with one pdz domain localized at cadherin-based cell-cell adhesion sites. The Journal of biological chemistry 1998;273: 3470–3475.

33. Kemmler W, Peterson JD, Steiner DF. Studies on the conversion of proinsulin to insulin. I. Conversion in vitro with trypsin and carboxypeptidase B. J Biol Chem 1971;246: 6786–6791.

34. Gupta R, Nguyen DC, Schaid MD, Lei X, Balamurugan AN, Wong GW, et al. Complement 1q-like-3 protein inhibits insulin secretion from pancreatic beta-cells via the cell adhesion G protein-coupled receptor BAI3. J Biol Chem 2018;293: 18086–18098.

35. Gregory B, Rahman N, Bommakanti A, Shamsuzzaman M, Thapa M, Lescure A, et al. Correction: The small and large ribosomal subunits depend on each other for stability and accumulation. Life Sci Alliance 2019;2.

36. Heijnen HF, van Wijk R, Pereboom TC, Goos YJ, Seinen CW, van Oirschot BA, et al. Ribosomal protein mutations induce autophagy through S6 kinase inhibition of the insulin pathway. PLoS Genet 2014;10: e1004371.

37. Borg J, Klint C, Wierup N, Strom K, Larsson S, Sundler F, et al. Perilipin is present in islets of Langerhans and protects against lipotoxicity when overexpressed in the beta-cell line INS-1. Endocrinology 2009;150: 3049–3057.

38. Mishra A, Liu S, Promes J, Harata M, Sivitz W, Fink B, et al. Perilipin 2 downregulation in beta cells impairs insulin secretion under nutritional stress and damages mitochondria. JCI Insight 2021;6.

39. Tsaytler P, Bertolotti A. Exploiting the selectivity of protein phosphatase 1 for pharmacological intervention. FEBS J 2013;280: 766–770.

