## Supplementary Information for "Mechanisms of spinophilin-dependent pancreas dysregulation underlying diabesity"

*Animals*

All animal studies were performed in accordance with the Guide for the Care and Use of Laboratory Animals and approved by the School of Science Institutional Animal Care and Use Committee (SC270R, SC310R) at Indiana University-Purdue University, Indianapolis (IUPUI). Male and Female $Lepr^{db/+}$ mice (B6.BKS(D)-Leprdb/J, Stock #000697, Jackson Laboratories, Bar Harbor, ME) and whole-body heterozygous spinophilin mice (Stock #018609; RRID: MMRRC_049172-UCD) were initially purchased from Jackson Laboratories and breeding colonies were maintained at IUPUI. Mice containing loxP sites around exon 3 of the *Ppp1r9b* (spinophilin) gene $Spino^{fl/fl}$ were generated by the University of Michigan and recently described

(23). Ins1Cre mice (B6(Cg)-Ins1tm1.1(cre)Thor/J; stock #026801) and Ins2$^{Akita}$ (C57Bl/6-Ins2$^{Akita}$/J; Stock #003548) mice were from Jackson laboratories. All animals were provided with chow and water *ad* libitum and group-housed. Mice were maintained on a 12-hour light/dark schedule (7 am-7 pm-7 am). Leptin receptor heterozygous mutant mice (Lepr$^{db/+}$) were crossed with heterozygous spinophilin mice (Spino$^{+/-}$) to generate Lepr$^{db/+}$/Spino$^{+/-}$ male and female breeders. These mice were crossed to generate the following genotypes that were used for the genetic obesity model studies: Lepr$^{+/+}$/Spino$^{+/+}$, Lepr$^{db/db}$/Spino$^{+/+}$, Lepr$^{+/+}$/Spino$^{-/-}$, and Lepr$^{db/db}$/Spino$^{-/-}$. Mice were weighed bi-weekly from 4 weeks until 20 weeks of age. These mice were provided with standard chow (LabDiet, St. Louis, MO, Diet #5001, 23% protein, 4.5% fat, 6% fiber) & water *ad libitum*. For HFF mice, spinophilin WT and spinophilin$^{-/-}$ male and female mice were weaned at Postnatal day (P) 21 and placed on high-fat diet (Research Diets, Inc., New Brunswick, NJ, D12492, 60% fat) at P28. These mice were weighed bi-weekly until 20 weeks of age. Spinophilin floxed mice were crossed with Ins1Cre mice to generate Spino$^{\Delta Ins}$ floxed line. These mice were placed on standard chow *ad* libitum after weaning at P21 and weighed bi-weekly until 18 weeks of age. These mice were sacked via decapitation without anesthesia at 20 weeks of age and pancreas was dissected, frozen in liquid nitrogen, and stored at -80ºC for further biochemical studies. No *a priori* sample size calculation was performed. Not all mice were measured every other week for weights, so the total number of animals per group was different at different weeks; however, an N of at least 4 per time point was used for all weights. For area under the curve, weights at 6, 8, and 10-weeks, and immunoblotting quantitation, each individual data point is shown.

*Immunoprecipitation from* WT, Ins2$^{Akita}$, or Lepr$^{db/db}$ mice.

Pancreatic tissue was dissected from WT, Ins2$^{Akita}$, or Lepr$^{db/db}$ mice. Tissue was homogenized in 2 mL of RIPA buffer (20 mM Tris HCl, 150 mM NaCl, 2 mM EDTA, 1X protease inhibitor cocktail (BiMake.com, Houston TX), phosphatase inhibitors (20 mM sodium fluoride, 20 mM sodium orthovanadate, 20 mM β-glycerophosphate, and 10 mM sodium pyrophosphate; MilliporeSigma, St. Louis, MO or Thermo-Fisher Scientific, Waltham MA), 1% NP-40 (Thermo-Fisher Scientific), 1% deoxycholate (Thermo-Fisher Scientific). Homogenates were sonicated and incubated with rotating for 1 hour at 4ºC. Homogenates were then centrifuged at 16,900 x g for 10 minutes at 4ºC. 10 µl of goat anti spinophilin antibody (Santa Cruz biotechnology, Dallas, TX, Catalog #14774 - discontinued) or 3 µl of rabbit anti spinophilin antibody (Cell Signaling Technologies, 9061S) were added to ~400 µl of supernatants. Antibody was incubated for 1 hr and then 20 µl of Protein G magnetic beads (Dynabeads, Life Technologies, Catalog #10009D) that had been washed 3 times in IP wash buffer (150 mM NaCl, 50 mM Tris-HCl pH 7.5, 0.5% (v/v) Triton X-100) were added. Beads were incubated for 1.5 hours and then washed 3 times in IP wash buffer. Beads were eluted in 40µl of 2X Laemmli sample buffer and 20 µl was run on a hand-cast SDS-PAGE gel for Coomassie staining and proteomics or immunoblotting.

*Coomassie Staining and tryptic digestion for Gel-C MS proteomics.*

SDS-PAGE gels containing spinophilin immunoprecipitates were stained with an Imperial colloidal Coomassie stain (ThermoFisher #24615) and regions of the gel were excised and tryptically digested as previously described[1, 2]. For all steps, sufficient volume of reagent was used that covered the gel pieces. Excised gels were destained (25 mM ammonium bicarbonate in 50 % acetonitrile (ACN)). DTT (10 mM) in 25 mM ammonium bicarbonate was added to reduce disulfides. Iodoacetamide (25 mM) was added to alkylate free-sulfhydryl groups and the reaction proceeded in the dark for 45 minutes. Gel pieces were subsequently incubated in 25 mM ammonium bicarbonate and then dehydrated with 25 mM ammonium bicarbonate in 50% ACN. The samples were then placed in a rotary vacuum and centrifuged until dry and subsequently digested with 12.5 ng/µl trypsin in 25 mM ammonium bicarbonate at 37ºC overnight. Supernatants were collected from all samples. Remaining gel pieces were washed with 5% formic acid in 50% ACN and were vortexed and sonicated for 5 minutes.

*Immunoblotting.*

Immunoblotting was performed as previously described[1, 2]. Briefly, SDS PAGE gels were transferred using either a wet or semi-dry transblot turbo apparatus (Bio-Rad, Hercules, CA). Immunoblots were probed with goat anti spinophilin antibody, mouse anti myosin-9 antibody (MilliporeSigma MABT164), mouse anti neurabin antibody (Santa Cruz Biotechnology, SC-136327), or mouse anti PP1$\alpha$ antibody (Santa Cruz Biotechnology, SC-7482) and infra-red secondary antibodies (From Jackson ImmunoResearch or Invitrogen) and developed on a LiCor Odyssey (LiCor Biosciences, Lincoln, NE).

*Proteomics for Gel-C MS.*

Tryptic digestions from a colloidal Coomassie-stained gel underwent proteomics analysis on a Q-Exactive mass spectrometer using higher-energy collisional dissociation as previously described [2, 3]. Specifically, Digested samples were loaded onto a 100 µm x 2 cm Acclaim PepMap100 C18 nano trap column (5 µm, 100 Å) (ThermoFisher) with an Ultimate 3000 liquid chromatograph (ThermoFisher) at 5 µL/min. The peptides were separated on a silica capillary column that was custom-packed with C18 reverse phase material (Magic, 0.075 mm x 150 mm, 5 µm, 120 Å,

Michrom Bioresources, Inc., Auburn, CA). The gradient was pumped at 300 nL/min from 10-45% solvent B (99.9% acetonitrile, 0.1% formic acid) for 87 min, then to 90% solvent B for 5 min, and re-equilibrated to solvent A (99.9% water, 0.1% formic acid) for 12 min. The mass spectrometer was operated in data-dependent acquisition mode controlled by the Xcalibur 2.2 software. Peptide mass spectra were acquired from an m/z range of 350-2000 at resolving power of 70,000 for 400 *m/z* ions. The top 15 most abundant multiply charged ions were subjected to higher-energy collisional dissociation (HCD) at a resolving power of 17,500 for 400 *m/z* ions. Ions with a charge state >+6 were rejected. AGC targets were set to 3e6 for MS1 and 1e5 for data-dependent MS2 with an underfill ratio of 2.5%, given an intensity threshold of 5.0e4. A dynamic exclusion of 10.0 s was used.

Data were searched in Proteome Discoverer using a SEQUEST plug-in (Version 1.4.1.14). The settings were: Peptide tolerance of 10.0 PPM (monoisotopic), Fragment Tolerance of 0.020 Da (monoisotopic), variable modifications +16 on Met (oxidation), +42 on Lys (Acetylation), +57 on Cys (carbamidomethylation), +80 on Ser, Thr, Tyr (Phosphorylation), +80 on Ser, Thr (Sulfation). There were no fixed modifications. Tryptic database was searched and up to 3 missed cleavages were permitted. Data were loaded into Scaffold and then exported to Excel. Supplemental tables are modified from the exported Excel table and the Scaffold file is included in the Supplemental Data (Scaffold_Supplement).

*Tandem Mass Tag proteomics of spinophilin immunoprecipitates.*

Sample preparation, mass spectrometry analysis, bioinformatics, and data evaluation for quantitative proteomics and phosphoproteomics experiments were performed in collaboration with the Indiana University School of Medicine Center for Proteome Analysis similarly to several previously published protocols [4, 5]. Specifically, spinophilin was immunoprecipitated as described above and Protein G magnetic beads were washed 3 times in PBS. After washing, beads were covered with 8 M Urea, 100mM Tris hydrochloride, pH 8.5, reduced with 5mM tris (2-carboxyethyl) phosphine hydrochloride (TCEP, Sigma-Aldrich Cat No: C4706) for 30 minutes at room temperature to reduce the disulfide bonds. The resulting free cysteine thiols were alkylated using 10 mM choloracetamide (CAA, Sigma Aldrich Cat No: C0267) for 30 minutes at RT, protected from light. Samples were diluted to 2 M Urea with 50 mM Tris pH 8.5 and proteolytic digestion was carried out with Trypsin/LysC Gold (0.3 µg, Mass Spectrometry grade, Promega Corporation Cat No: V5072) overnight at 35 °C. After digestion, samples were quenched with 0.4% trifluoroacetic acid (v/v, Fluka Cat No: 91699) and the resultant peptides were desalted by solid phase extraction using C18 Spin columns (Pierce Cat. No. 89870).

Peptides were reconstituted in 20 µL of 50 mM triethylammonium bicarbonate (TEAB, Sigma-Aldrich Cat No: T7408), pH 8.5 and labeled with 0.20 mg aliquots of TMT10plex™ Isobaric Label Reagent (Thermo Fisher Scientific Cat No: 90111, Lot WG320953, Table 1). After one hour incubation, the labeling reaction was quenched with 0.3% hydroxylamine (final v/v) for 15 minutes before combining the samples. The multiplexed sample was concentrated to dryness in vacuum centrifuge, reconstituted with 0.1% TFA aq. (v/v), desalted via Waters Sep-Pak® Vac cartridge, and speed vacced to dryness. 1/10$^{th}$ of the total sample was then injected using an Easynano LC1200 coupled with 25cm Aurora column (Ionopticks AUR2-25075C18A) on an Eclipse Orbitrap mass spectrometer (Thermo Fisher Scientific). Peptides were eluted over a 180-minute method: Solvent B was increased from 5%-30% over 160 min, to 85% Bover 10 min, and down to 10% B (Solvent A: water, 0.1% formic acid; Solvent B: 100% acetonitrile, 0.1% formic acid). The mass spectrometer was operated in positive ion mode with 3 FAIMS CVs (-45, -55, -65). A cycle time of 1 s was used for each CV. MS1 parameters for each cycle were: orbitrap resolution of 120,000, scan range of 350-1600 m/z, standard AGC, 50 ms max IT, minimum intensity of 2.5e4, precursor fit of 70% 0.7 m/z, charge state 2-6, 60 sec dynamic exclusion. MS2 settings were quadrupole isolation of 0.7 m/z, fixed HCD of 34, orbitrap resolution of 50,000, 200% AGC, dynamic max IT.

Data were analyzed in Proteome Discoverer 2.5. A M*us musculus* protein database (UniProtKB/TrEMBL; last modified Jan 09, 2017) plus common laboratory contaminants was searched using SEQUEST HT. Precursor mass tolerance was set to 10 ppm and fragment mass tolerance set at 0.02 Da with a maximum of 3 missed cleavages. Dynamic modifications include methionine oxidation; deamidation of asparagine, phosphorylation on serine, threonine, and tyrosine, TMT on lysine, acetyl on lysine, and GG+TMT (+343.206) on lysine residues. Dynamic peptide modifications were TMT at the N-terminus; dynamic protein terminus modifications were acetylation, met-loss, and met-loss plus acetylation. Static modifications were carbamidomethylation on cysteines. IMP-ptmRS node was used for localization scoring. Percolator false discovery rate (FDR) filtration of 1% was applied to both the peptide-spectrum match and protein levels. For the Proteome Discoverer consensus workflow, isobaric impurities corrections were turned on, the reporter ion co-isolation threshold was set to 50%, and the average signal to noise threshold was 5. All peptides were used for protein roll-up, but modified peptides were excluded for pairwise ratio testing. Proteomics data files are uploaded on the MassIVE repository (accession MSV000091159) and will be accessible upon publication.

For analyses of WT/KO log2-fold change, one KO-HFD sample was excluded from the calculation due to a high abundance (TMT quantitation) of spinophilin in the sample, suggesting either carry-over or a mis-genotyping (e.g. heterozygous animal). In addition, one Chow-WT sample showed excessively high levels of cytoskeletal proteins, potentially suggesting non-specific binding (e.g. levels 4-5 times higher than any other sample). Both the HFD-KO and the Chow-WT are shown in the supplemental table, but calculations for the tables in the text and for the stringdb were performed without these samples included. In addition, **Table S5** shows all protein abundances detected in the mass spectrometry run. This includes contaminants.

| **Sample ID** | **TMT label** |
|---|---|
| WT_HFD | 126 |
| WT_HFD | 127N |
| WT_HFD | 128N |
| WT_StdChw | 128C |
| WT_StdChw | 129N |
| WT_StdChw | 129C |
| spKO_HFD | 130N |
| spKO_HFD | 130C |
| spKO_HFD | 131 |

**Figure S1. Comparison of Controls. A-C.** AUC was compared between floxed control mice and INS1Cre control mice at 6,8, and 10 weeks to determine any significant difference. We found no significant difference in AUC when we compared the different controls in our GTT and ITT (normalized and not normalized to baseline) experiments. **A.** $t(9)=0.3858$, $p=0.7086$. **B.** $t(9)=0.1843$ $p=0.8579$. **C.** $t(5)=.1471$, $p=0.8888$. **D.** $t(5)=0.4887$, $p=0.6458$.

**Figure S2. ITT graph A.** ITT of 8 week-old Spino$^{\Delta Ins}$ & controls analyzed from raw glucose values, given in mg/dL $(F(2,10))=0.8438$, $p=0.4585$. **B.** Area under curve for ITT of 8-week-old Spino$^{\Delta Ins}$ & controls. An unpaired t-test analyzing the AUC revealed no significant difference in insulin tolerance $t(11)=1.271$, $p=0.2301$.

### Figure S1

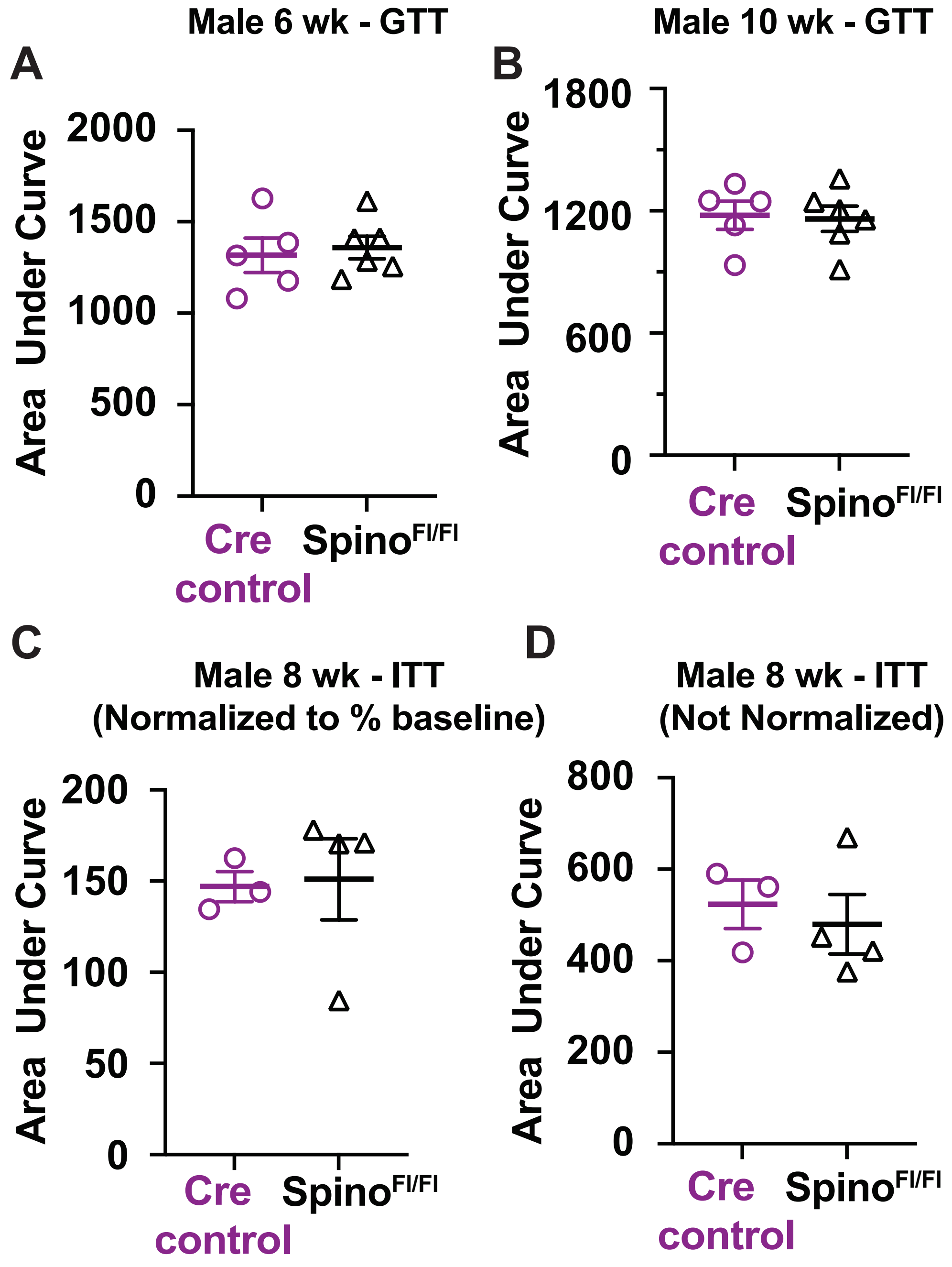
A
Male 6 wk - GTT
Area Under Curve
2000
1500
1000
500
0
Cre control
Spino$^{Fl/Fl}$
B
Male 10 wk - GTT
Area Under Curve
1800
1200
600
0
Cre control
Spino$^{Fl/Fl}$
C
Male 8 wk - ITT
(Normalized to % baseline)
Area Under Curve
200
150
100
50
0
Cre control
Spino$^{Fl/Fl}$
D
Male 8 wk - ITT
(Not Normalized)
Area Under Curve
800
600
400
200
0
Cre control
Spino$^{Fl/Fl}$

Figure S2

Male 8 wks old

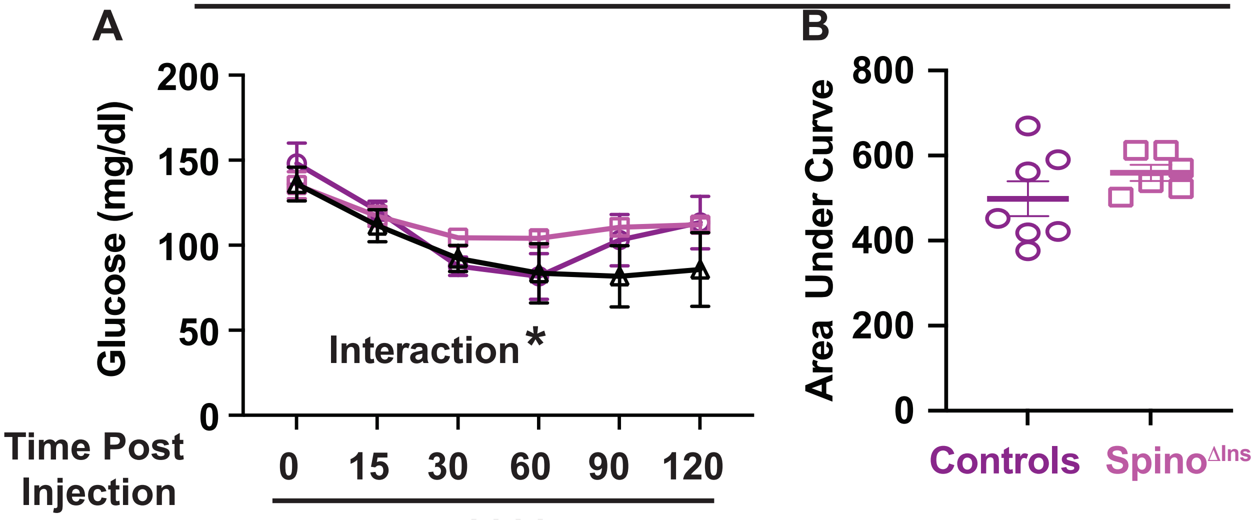

1. Edler, M.C., et al., *Mechanisms Regulating the Association of Protein Phosphatase 1 with Spinophilin and Neurabin.* ACS Chem Neurosci, 2018. **9**(11): p. 2701-2712.
2. Salek, A.B., et al., *Spinophilin regulates phosphorylation and interactions of the GluN2B subunit of the N-methyl-d-aspartate receptor.* J Neurochem, 2019. **151**(2): p. 185-203.
3. Hiday, A.C., et al., *Mechanisms and Consequences of Dopamine Depletion-Induced Attenuation of the Spinophilin/Neurofilament Medium Interaction.* Neural Plast, 2017. **2017**: p. 4153076.
4. Mosley, A.L., et al., *Highly reproducible label free quantitative proteomic analysis of RNA polymerase complexes.* Mol Cell Proteomics, 2011. **10**(2): p. M110 000687.
5. Grecco, G.G., et al., *A multi-omic analysis of the dorsal striatum in an animal model of divergent genetic risk for alcohol use disorder.* J Neurochem, 2021. **157**(4): p. 1013-1031.
